# An infant mouse model of influenza virus transmission demonstrates the role of virus-specific shedding, humoral immunity, and sialidase expression by colonizing *Streptococcus pneumoniae*

**DOI:** 10.1101/456665

**Authors:** Mila Brum Ortigoza, Simone Blaser, M. Ammar Zafar, Alexandria Hammond, Jeffrey N. Weiser

## Abstract

The pandemic potential of influenza A viruses (IAV) depends on the infectivity of the host, transmissibility of the virus, and susceptibility of the recipient. While virus traits supporting IAV transmission have been studied in detail using ferret and guinea pig models, there is limited understanding of host traits determining transmissibility and susceptibility because current animal models of transmission are not sufficiently tractable. Although mice remain the primary model to study IAV immunity and pathogenesis, the efficiency of IAV transmission in adult mice has been inconsistent. Here we describe an infant mouse model which support efficient transmission of IAV. We demonstrate that transmission in this model requires young age, close contact, shedding of virus particles from the upper respiratory tract (URT) of infected pups, the use of a transmissible virus strain, and a susceptible recipient. We characterize shedding as a marker of infectiousness that predicts the efficiency of transmission among different influenza virus strains. We also demonstrate that transmissibility and susceptibility to IAV can be inhibited by humoral immunity via maternal-infant transfer of IAV-specific immunoglobulins, and modifications to the URT milieu, via sialidase activity of colonizing *Streptococcus pneumoniae* (Spn). Due to its simplicity and efficiency, this model can be used to dissect the host’s contribution to IAV transmission and explore new methods to limit contagion.

**IMPORTANCE:** This study provides insight into the role of the virus strain, age, immunity, and URT flora on IAV shedding and transmission efficiency. Using the infant mouse model, we found that: (a) differences in viral shedding of various IAV strains is dependent on specific hemagglutinin (HA) and/or neuraminidase (NA) proteins; (b) host age plays a key role in the efficiency of IAV transmission; (c) levels of IAV-specific immunoglobulins are necessary to limit infectiousness, transmission, and susceptibility to IAV; and (d) expression of sialidases by colonizing Spn antagonize transmission by limiting the acquisition of IAV in recipient hosts. Our findings highlight the need for strategies that limit IAV shedding, and the importance of understanding the function of the URT bacterial composition in IAV transmission. This work reinforces the significance of a tractable animal model to study both viral and host traits affecting IAV contagion, and its potential for optimizing vaccines and therapeutics that target disease spread.

## INTRODUCTION

Influenza virus infections continue to cause 140,000-700,000 hospitalizations and 12,000-56,000 deaths in the United States annually (1). For the 2017-2018 season alone, more than 900,000 people were hospitalized and 80,000 people died from influenza (2). Despite the availability of vaccines which have been efficacious at preventing hospitalizations, morbidity, and mortality, evidence that the inactivated influenza virus (IIV) vaccine blocks virus acquisition, shedding, or transmission has been limited in animal models (3-7). In addition, the low vaccination coverage (in the population) and low vaccine effectiveness (due to viral antigenic drift) likely contributes to the limited effects of the IIV vaccine (8, 9). Likewise, available therapeutics, primarily neuraminidase inhibitors (NI), have been shown to be effective at reducing the duration of illness if treatment is initiated within 24 hours of symptom onset (10-13). However, NI treatment of index cases alone shows limited effectiveness reducing viral shedding or transmission, possibly due to its short therapeutic window (10, 11, 14, 15). These limitations of our current options to prevent disease spread highlight a critical aspect of the IAV ecology that needs further study: contagion.

While IAV transmission has been studied in human, ferret, and guinea pig models, there is a general lack of understanding about the host influence on viral transmission, because none of these models are easily manipulated. Hence, scientific progress to date has emphasized viral genetics, viral tropism, and environmental impacts on transmission (16-19). While these factors contribute to knowledge about IAV contagion, host characteristics that could affect transmissibility, including the highly variable composition of the URT flora, remain largely unexplored.

This knowledge gap could be addressed with the use of mice, whose practical features (small, inexpensive, inbred), expansive reagent repertoire, and availability of genetically-modified hosts allows for studies of extraordinary intricacy providing a significant research advantage. Since the 1930s, the mouse model has been essential in understanding IAV immunity and pathogenesis, and early studies described its usefulness in evaluating IAV transmission (20, 21). However, the use of mice for studying IAV transmission has been largely disregarded due to marked differences among studies and low transmission rates (22-24). Nevertheless, recent reports have revived the potential of the murine species as an IAV transmission model (23-28). Therefore, in this study, we sought to reevaluate the mouse as a tool to study the biology of IAV contagion, particularly the contribution of host factors.

## RESULTS

### Infant mice support efficient influenza virus transmission

Given the remarkable capacity of infant mice to support IAV transmission among littermates (25), we sought to validate and optimize the infant mouse as a potential new model to study IAV transmission. Restricted URT infection of infant C57BL/6J pups in a litter (index) was performed with low volume intranasal (IN) inoculum (3μl) using IAV strain A/X-31 (H3N2) (24, 29). Intralitter transmission was assessed in littermates (contact/recipient) by measuring virus from retrograde tracheal lavages at 4-5 days post-infection (p.i.) (Fig. 1A). A/X-31 virus was selected to model transmission because of its intermediate virulence in mice (30) and ability to replicate in the URT to high titers with natural progression to the lungs, simulating key features of the infectious course in humans. Furthermore, the 50% mouse infectious dose (MID_50_) in this model is 4-5 plaque forming units (PFU), suggesting high susceptibility to A/X-31 infection. Transmission efficiency was observed to be 100% when index and contact pups were housed together at the time of IAV inoculation (Fig. 1B). Transmission declined the longer the index and recipient pups were housed apart prior to being in direct contact, and was completely eliminated when housed together after 72 hours of separation (Fig. 1C and Fig. S1). This observation suggested that in this model, transmission from index to recipient is most effective within the first 72-hour period of contact.

**Figure 1.**
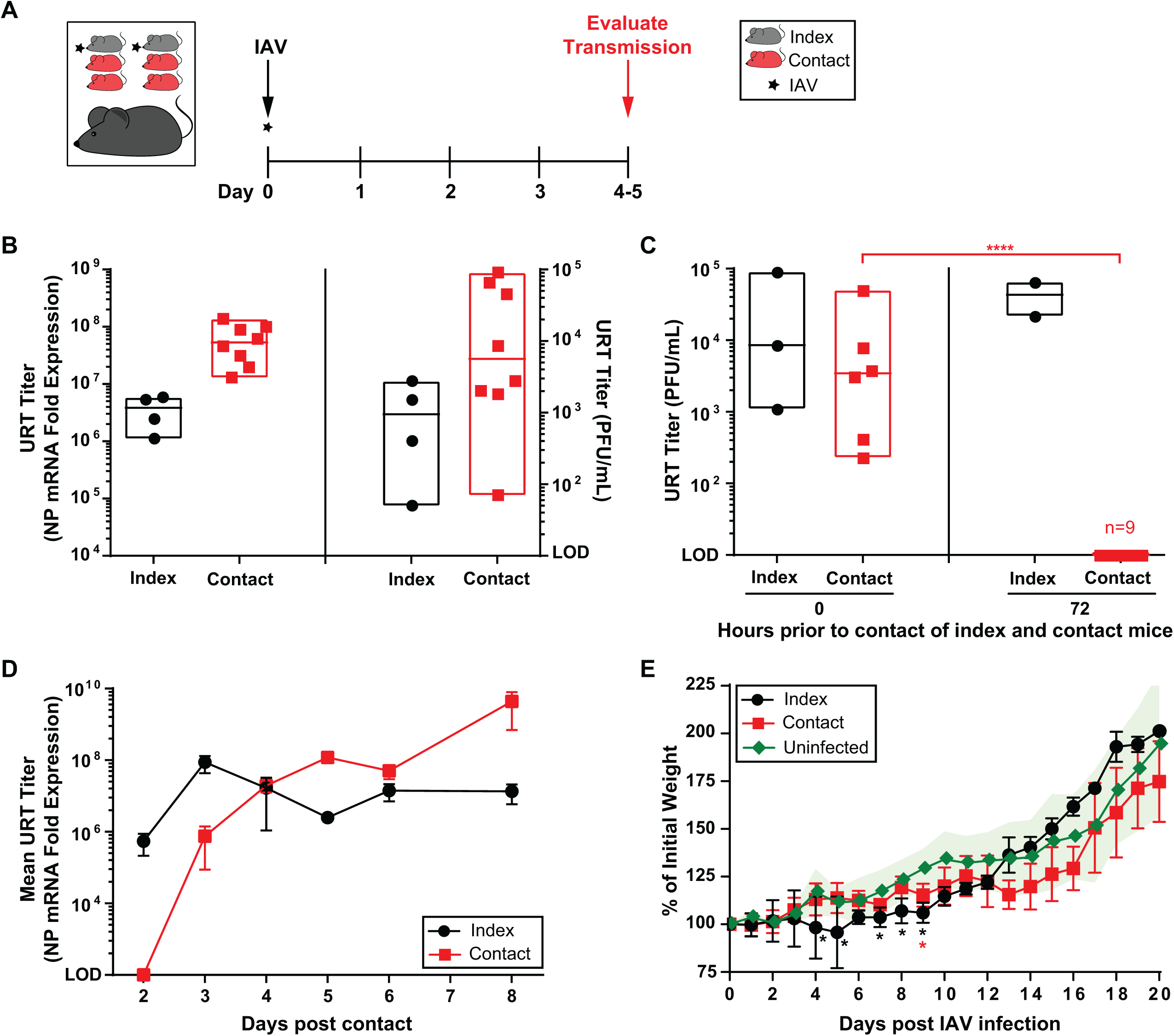
Transmission of IAV in infant mice. (A) Schematic and timeline of experimental design. Index and contact pups were arbitrarily assigned, maintained in the same cage, and cared for by the same mother. At day 0 (4-7 days of age), pups were infected IN with 250 PFU of A/X-31 (index), and cohoused with uninfected littermates (contact) for 4-5 days prior to evaluating for transmission. (B) Transmission of IAV to contact pups was evaluated via qRT-PCR (left panel) or plaque assay (right panel) from retrograde URT lavages after sacrifice. URT titers are represented by a box plot extending from the minimum to maximum values for each data set. Each symbol represent the titer measured from a single pup with median values indicated by a line within the box. Index and contact pups are shown in black and red symbols, respectively. (C) Window of transmission was evaluated by separating index and contact pups for a defined period prior to contact. After infection of index pups, uninfected contact pups were housed apart (in a separate cage) for 0 and 72 hours prior to cohousing with infected index for 5 days. Transmission to contact pups was evaluated via plaque assay from retrograde URT lavages. URT titers are represented by a box plot as described above. (D) Timecourse of A/X-31 transmission. Pups in a litter were subjected to an A/X-31 transmission experiment (described above) and transmission to contact pups was evaluated via qRT-PCR from retrograde URT lavages at indicated day post contact. Mean URT titers ± standard deviation (SD) are represented. (E) Morbidity of A/X-31 infection in index and contact pups over the course of 20 days. Pups in a litter were subjected to an A/X-31 transmission experiment (described above) and weight of each pup was measured daily. Percent of initial weight ± SD is represented (uninfected group n=9, index group n=3-4, contact group n=4-5). Differences among group means were analyzed using the Student’s *t* test. All panels represent at least two independent experiments. * *p*<0.05, ** *p*<0.01, IAV=Influenza A virus, URT=upper respiratory tract, NP=nucleoprotein, PFU=plaque forming unit, LOD=Limit of detection.

To determine the window of viral acquisition in recipient mice, an 8-day IAV transmission experiment was performed (Fig. 1D). The observed growth of IAV in the URT of recipient pups suggested that *de novo* virus acquisition occurred between 2-3 days after contact with the index. Hence, the infectious window for the index pups corresponded with the timing of IAV acquisition in contact pups.

Given that pups gain weight as they grow, morbidity in this model was assessed by observing a decrease in weight gain during the infectious period. Mild morbidity of pups was observed in both the index and contact groups, with complete recovery from IAV infection by 10 days p.i. (Fig. 1E).

### Direct contact between pups is required for influenza virus transmission

Because infant mice need their mother for survival during the first 21 days of life, they cannot be separately housed, therefore this model cannot differentiate between airborne versus droplet routes of transmission. To distinguish between direct and indirect contact routes of transmission, the mother and housing contents were evaluated as potential fomites. This was done by daily switching the mothers or the cages with bedding between infected and uninfected litters, respectively (Fig. S2A-B). Inefficient or no transmission was observed, suggesting that direct (close) contact between pups is the main mode of transmission. Occasionally during a transmission experiment, the mother in the cage became infected with IAV from close contact with her infected pups (Fig. S2C). Although the acquisition of IAV in the mother was a rare event, we did not observe a decline in transmission in contact pups when the mother did not become infected, despite the mother being capable of transmitting IAV to her pups if she were to be inoculated with IAV as the index case (Fig. 2D).

**Figure 2.**
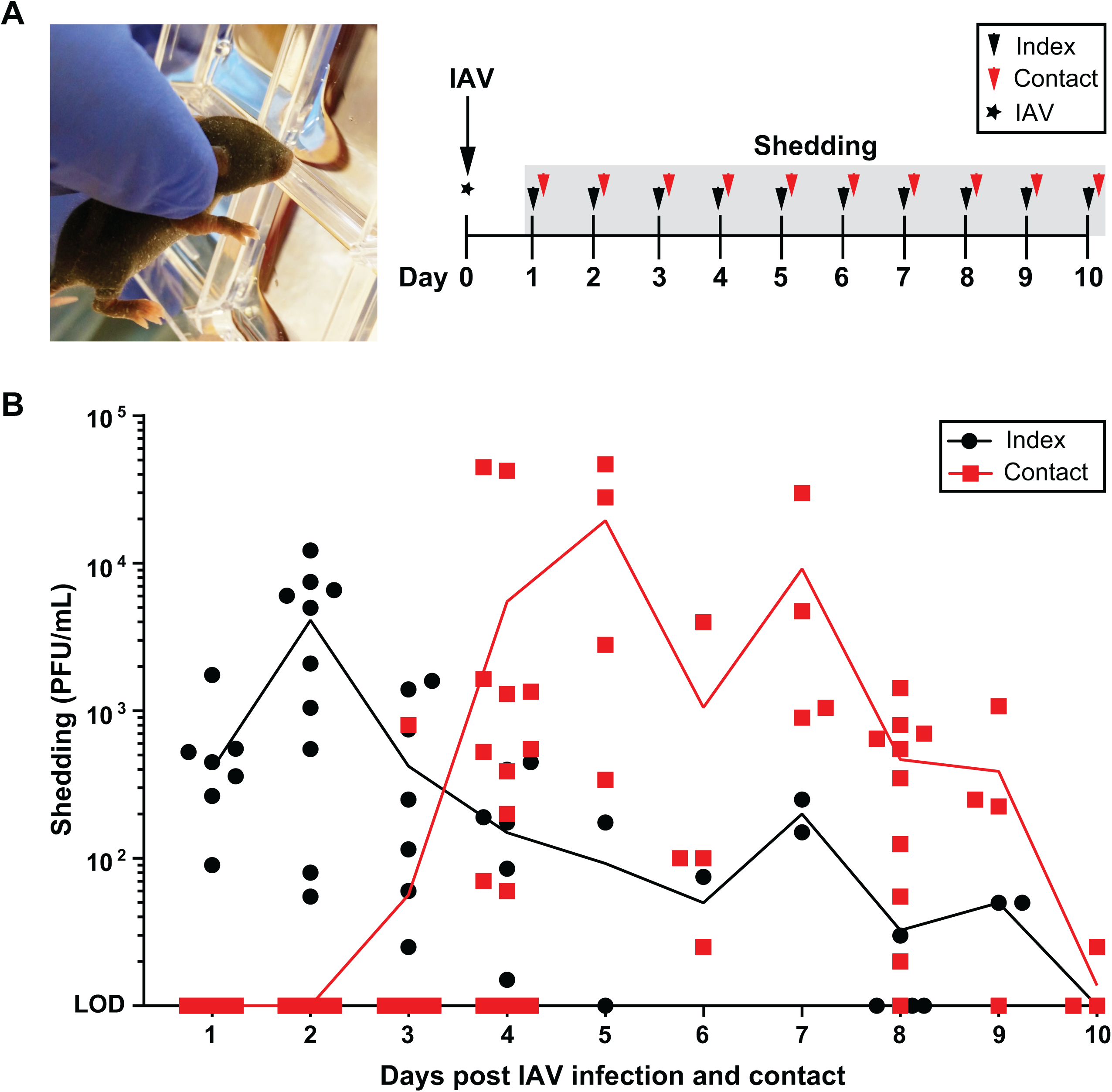
Shedding of IAV. (A) Image of infant mouse shedding procedure and schematic timeline of experimental design. At day 0 (4-7 days of age), pups were infected IN with 250 PFU of A/X-31 (index), and cohoused with uninfected littermates (contact) for 10 days. Shedding of IAV was collected by dipping the nares of each mouse in viral media daily. (B) Shedding samples from each day were evaluated individually via plaque assay. Each symbol represents the shedding titer measured from a single mouse at the specified day. Index and contact pups are shown in black and red symbols, respectively. Mean values are connected by a line. IAV=Influenza A virus, PFU=plaque forming unit, LOD=Limit of detection.

### Role of shedding of influenza virus from the upper respiratory tract

To determine the correlates of transmission, an assay was developed to quantify infectious virus expelled from the nasal secretions of pups. This assay allowed us to follow the journey of particle exit from index pups to acquisition by contact pups over the course of the transmission period. Index pups in a litter were infected with A/X-31 and cohoused with uninfected littermates for 10 days. The nares of each mouse was gently dipped in viral media daily, and virus titers were assessed for each sample (Fig. 3A). We observed that index pups, like in humans (31), began shedding virus from day 1 p.i., whereas recipient pups, who acquired IAV infection between day 2-3 (Fig. 1D), began shedding virus from day 4 post contact (Fig. 3B). This pattern of virus transit suggested that the timing of peak shedding from the index (days 1-3) corresponded with the timing of transmission to recipient pups (days 2-4) (Fig. S3) further confirming that a key determinant of IAV transmission in this model is shedding of virus from the secretions of index pups. Notably, detectable shedding in the contacts lagged transmission (higher transmission rate compared to number of contacts shedding virus), because of the period of viral replication required prior to the detection of shed virus (Fig. S3).

**Figure 3.**
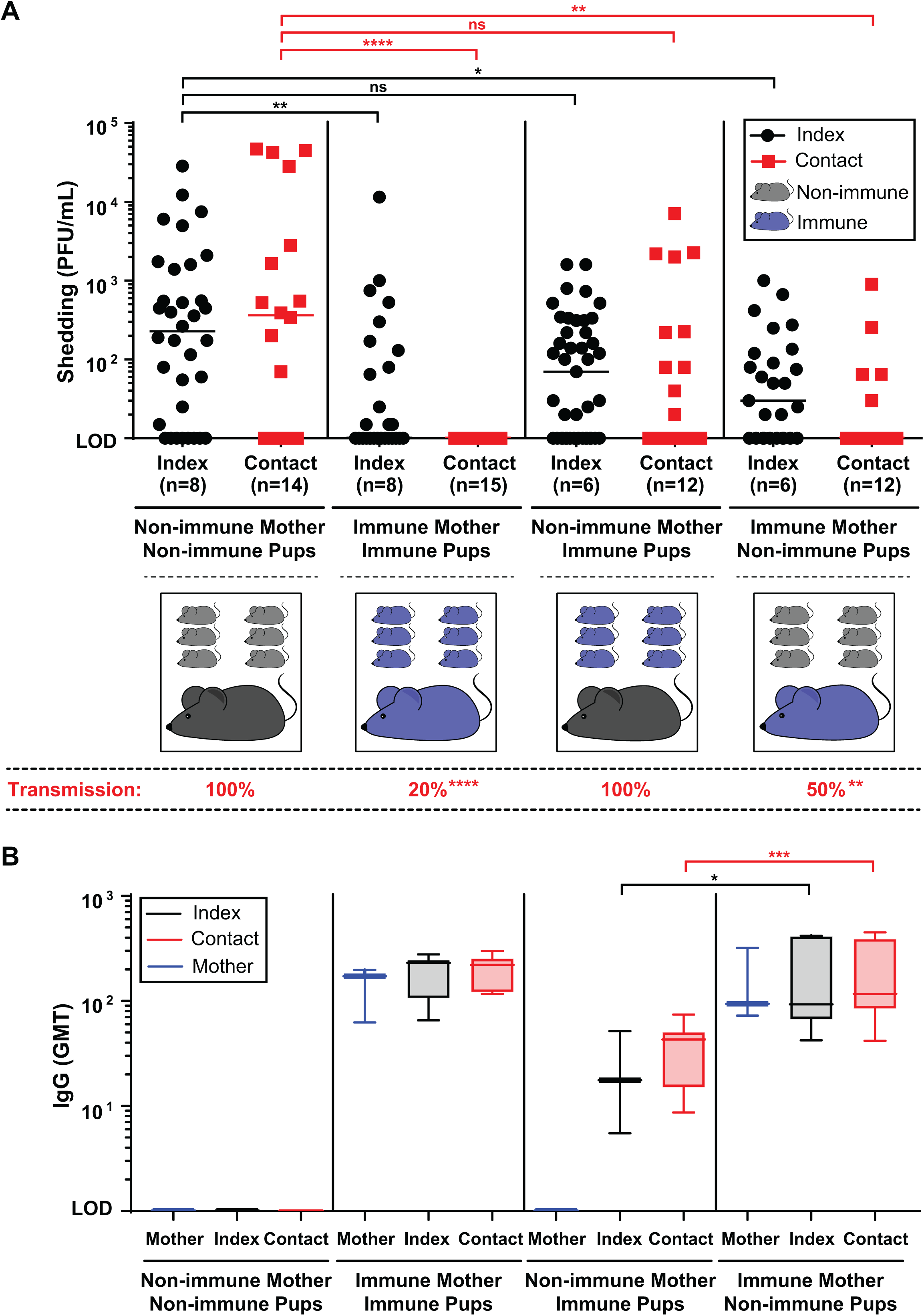
Maternal-infant transfer of IAV-specific immunity limits IAV shedding and transmission. (A) Adult females were infected IN with 250 PFU of A/X-31 and were left to recover from infection prior to breeding. Soon after birth, pups born from previously infected (immune) mothers or those born from non-immune mothers were either left with their biological mother or exchanged with a foster mother of opposite immune status. Pups paired with their biological or foster mothers were left to acclimate until 4-5 days of life, prior to being subjected to an IAV transmission experiment. Schematic for each experimental condition is shown. Pups in a litter were infected IN with 250 PFU of A/X-31 (index), and cohoused with uninfected littermates (contact) for 5 days. Shedding of IAV was collected by gently dipping the nares of each mouse in viral media daily. Shedding samples from each day were evaluated individually via plaque assay for each pup. Shedding titers shown represent pooled values for days 1-5 for index pups and days 4-5 for contact pups, representing days of maximum shedding for each group (as per Fig. 2). Each symbol represent the shedding titer measured from a single mouse for a specific day. Index and contact pups are shown in black and red symbols, respectively. Median values are indicated. At the end of 5 days, pups and mothers were sacrificed, and transmission to contact pups was evaluated via plaque assay from retrograde URT lavages. Percentage of transmission among contact pups is displayed below the graph. (B) Serum from mother and pups were obtained at the time of sacrifice. Samples from individual mice were evaluated for IAV-specific IgG by ELISA. IgG geometric mean titers (GMT) are represented by a box plot extending from the 25^th^ to 75^th^ percentile for each data set. Whiskers for each box encompasses the minimum to maximum values. Median values are indicated by a line within the box. All panels represent at least two independent experiments. Differences in transmission were analyzed using the Fisher’s exact test. * *p*<0.05, ** *p*<0.01, *** *p*<0.001, **** *p*<0.0001, PFU=plaque forming unit, LOD=Limit of detection, GMT=Geometric mean titer.

### Transmission efficiency of influenza viruses in mice is virus and age-dependent

Virus strain has been shown to be important in the efficiency of transmission in adult mice (20, 23, 26, 32). We thus tested the capacity of infant mice to support transmission of other IAV subtypes and an influenza B virus (IBV) (Table 1). Transmission among pups was greater for influenza A/H3N2 viruses and IBV, but lower for A/H1N1 viruses. Notably, A/X-31 was more efficiently transmitted compared to its parent A/PR/8/1934 virus, suggesting that either the HA and/or NA proteins are responsible for efficient shedding and transmission of IAV in infant mice.

**TABLE 1.**
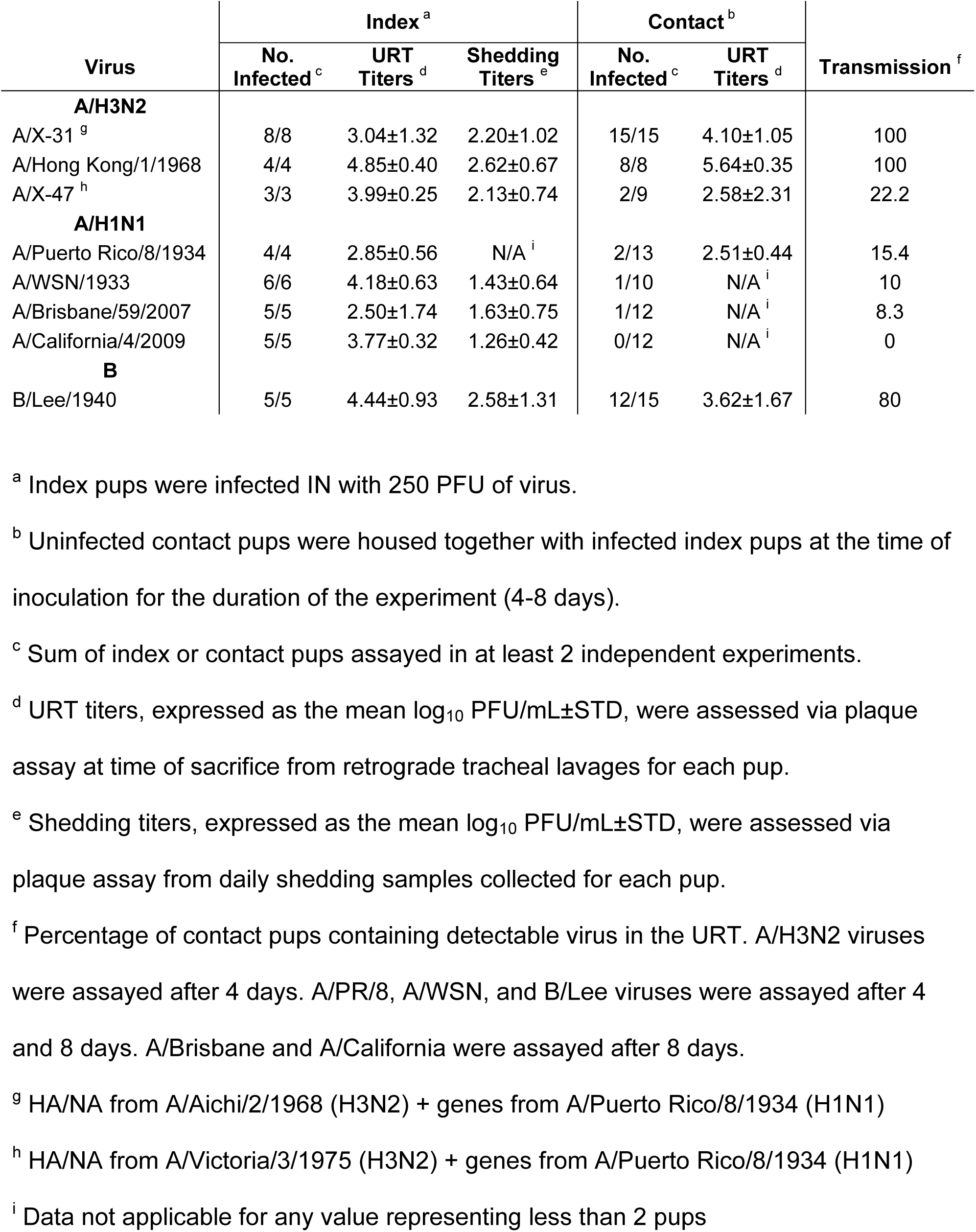
Transmissibility of influenza viruses in infant mice.

Surprisingly, the mean viral URT titers in index pups did not correlate with IAV transmission (*r*=0.315), indicating that virus replication in the URT alone was insufficient to mediate effective transmission. To determine the cause of the differences in transmission efficiencies observed among virus strains, the shedding for each virus was analyzed. We observed that virus shed from index pups correlated with IAV transmission (*r*=0.8663), further supporting virus shedding as the main determinant of IAV transmission efficiency in infant mice (Fig. S4A-B).

Given the effectiveness of the infant mouse in supporting transmission of IAV, we evaluated the disparities of transmission efficiency previously reported in adult mice (20-23, 26). Mice infected with A/X-31 at different ages were housed with uninfected age-matched contacts, and transmission efficiency was assessed at 5 days p.i. (Table 2). We observed that 100% transmission was sustained in mice up to 7-days of age. Weaned and active adult mice (>28 days of age) failed to sustain efficient transmission altogether. Furthermore, mouse age correlated with transmission rate among contact mice (*r*=-0.8346) (Fig. S4C), confirming that in the murine model, the requirement for young age is necessary to support efficient IAV transmission. Although the transmission experiments in this study were done with an IAV inoculum of 250 PFU, and increasing the inoculum size to 10^3^-10^5^ PFU correlated with increasing IAV titers in the URT tract of index mice (*r*=0.9264), inoculum size was not associated with more efficient transmission among adult mice (*r*=-0.2582) (Fig. S4D).

**TABLE 2.**

| Mice Age <sup>c</sup> | Index <sup>a</sup> |  | Contact <sup>b</sup> |  | Transmission <sup>f</sup> |
| --- | --- | --- | --- | --- | --- |
|  | No. Infected <sup>d</sup> | URT Titers <sup>e</sup> | No. Infected <sup>d</sup> | URT Titers <sup>e</sup> |  |
| 4 | 3/3 | 4.17±0.77 | 4/4 | 4.66±0.50 | 100 |
| 7 | 7/7 | 4.40±1.23 | 11/11 | 3.94±1.04 | 100 |
| 14 | 9/9 | 3.12±0.95 | 11/17 | 3.41±1.00 | 64.7 |
| 21 | 4/4 | 3.21±0.78 | 6/9 | 2.82±0.78 | 66.7 |
| **Weaned <sup>g</sup>** |  |  |  |  |  |
| 28 | 3/3 | 3.80±0.86 | 0/6 | N/A <sup>h</sup> | 0 |
| 35 | 5/5 | 3.15±0.52 | 0/8 | N/A <sup>h</sup> | 0 |
| 56 | 6/6 | 2.50±0.71 | 1/9 | N/A <sup>h</sup> | 11.1 |

<sup>a</sup> Index mice were infected IN with 250 PFU of virus.
<sup>b</sup> Uninfected age-matched contact mice were housed together with infected index mice at the time of inoculation for the duration of the experiment (5 days).
<sup>c</sup> Age of mice expressed in days after birth.
<sup>d</sup> Sum of index or contact mice assayed in at least 2 independent experiments.
<sup>e</sup> URT titers, expressed as the mean log<sub>10</sub> PFU/mL±STD, were assessed via plaque assay at time of sacrifice from retrograde tracheal lavages for each mice.
<sup>f</sup> Percentage of contact mice containing detectable virus in URT after 5 days of contact.
<sup>g</sup> Mice weaned from breastfeeding and separated from the mother.
<sup>h</sup> Data not applicable for any value representing less than 2 pups

### Humoral immunity from prior influenza virus infection limits shedding and transmission

To further validate the relationship of viral shedding and transmission, we evaluated the role of IAV-specific immunity in this model. Because pups are infected at a young age and lack a fully functional adaptive immune response, it was necessary to provide IAV immunity via the mother (from prior IAV infection), who would then transfer immunoglobulins to her pups either pre-natally via trans-placental passage or post-natally via breastfeeding. Pups from immune mothers who were subjected to an intralitter IAV transmission experiment shed significantly fewer virus over the first 5 days of infection compared to pups from non-immune mothers (Fig. 3A left). Reduced shedding was associated with decreased transmission (20%) among immune litters (*p*<0.0001) (Fig. 3A below graph). To determine if the passage of anti-IAV immunoglobulins occurred pre-natally or post-natally, mothers were switched shortly after delivery such that an immune mother raised pups from a non-immune mother or vice-versa. These cross-foster experiments demonstrated that maternal passage of immunoglobulins either pre-natally or post-natally decreased IAV shedding amongst all pups, and that transmission to contact pups was more efficiently blocked when maternal antibodies were passed post-natally via breastfeeding (*p*<0.01) (Fig. 3A right). IAV-specific serum IgG was detected in immune mothers and pups born or cared by immune mothers, with the transfer of IgG via breastfeeding yielding higher titer of antibodies in these pups (Fig. 3B). IAV-specific serum IgA was detected in previously-infected mothers but unlike IgG was not passed to their pups in significant amounts (Fig. S4).

### *Streptococcus pneumoniae* colonization of the upper respiratory tract decreases influenza virus acquisition via bacterial sialidase activity

There is increasing evidence of the important role of the host’s gut microbiome on IAV-specific immunity in the respiratory tract (33, 34). Yet, there is only one study evaluating the role of the URT microbiota in IAV infection (35) and no studies on its effect on transmission. This is surprising given that the nasopharynx, a non-sterile environment extensively colonized by a diverse bacterial flora, is the first location encountered by IAV. Since Spn carriage is highest in children (36, 37), Spn colonization often precedes IAV infection in childhood (38, 39). Given that infant mice support efficient Spn colonization in the URT (25, 40), we investigated the impact of Spn colonization on IAV transmission. All pups in a litter were colonized with Spn prior to IAV infection of index pups to control for the efficient pup-to-pup transmission of Spn in the setting of IAV infection (25). IAV shedding was collected daily for each pup prior to evaluating for IAV transmission in contact littermates at 4 days p.i (Fig. 4A). We observed that the Spn colonized contact mice acquired IAV at a decreased rate (32%) as compared to uncolonized mice (100%) (*p*<0.0001) (Fig. 5B below graph), which corresponded to lower viral shedding among colonized contacts (Fig. 5B left). Since index Spn colonized and uncolonized mice infected with IAV (via inoculation) shed IAV at similar levels, this suggested an antagonistic effect of Spn colonization on IAV transmission through decreased acquisition by contact mice (Fig. 5B left).

**Figure 4.**
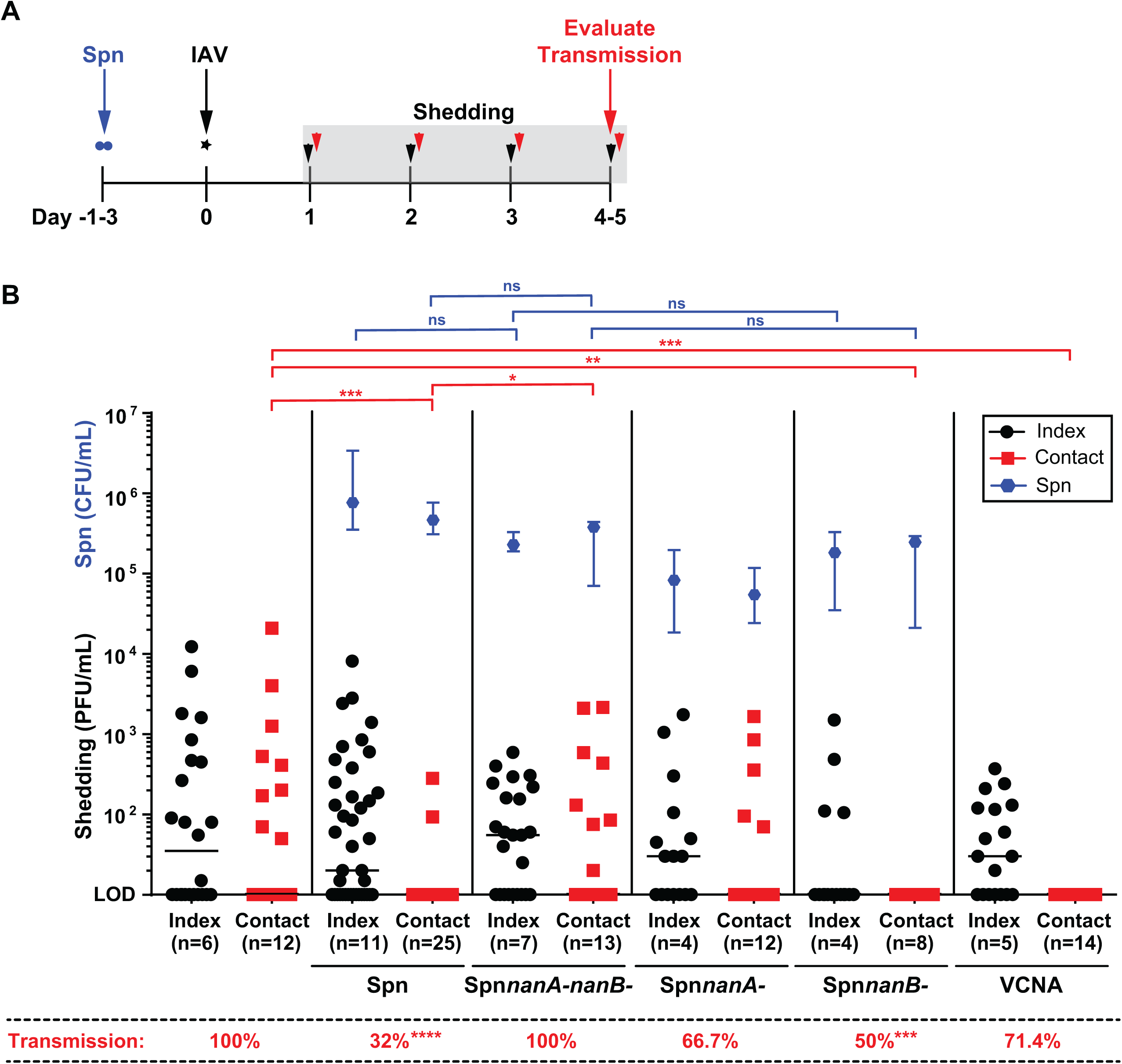
*Streptococcus pneumoniae* sialidases limit acquisition of IAV via transmission. (A) Schematic timeline of experimental design. At day -1 or -3 (3-4 days of age), all pups in a litter were colonized IN with either wild-type Spn; mutant Spn lacking NanA, NanB, or both; or treated IN with *Vibrio cholerae* neuraminidase (VCNA) twice daily. At day 0, pups were infected IN with 250 PFU of A/X-31 (index), and cohoused with uninfected littermates (contact) for 4 days. Shedding of IAV was collected by dipping the nares of each mouse in viral media daily. Transmission to contact pups was evaluated at day 4. (B) Shedding samples from each day were evaluated individually via plaque assay for each pup. Shedding titers shown represent pooled values for days 1-4 for index pups and days 3-4 for contact pups. Each symbol represent the shedding titer measured from a single mouse for a specific day with median values indicated. Index and contact pups are shown in black and red symbols, respectively. At the end of 4 days, pups were sacrificed, and transmission to contacts was evaluated via plaque assay from retrograde URT lavages. Percentage of transmission among contact pups is displayed below the graph. Density of colonizing Spn was measured in URT lavage samples of each pup. Each blue symbol represents the median Spn density ± interquartile range for each group. Differences in transmission were analyzed using the Fisher’s exact test. * *p*<0.05, ** *p*<0.01, *** *p*<0.001, **** *p*<0.0001, PFU=plaque forming unit, CFU=colony forming unit, LOD=Limit of detection.

Previous studies showed that sialidases expressed by colonizing Spn depletes host sialic acid (SA) from the epithelial surface of the murine URT, allowing Spn to utilize free SA for its nutritional requirements (41). Given that IAV requires SA for efficient attachment, we evaluated the role of Spn sialidases on IAV acquisition. We generated a double mutant lacking two common Spn sialidases: NanA and NanB (Spn*nanA^-^nanB^-^*), and tested its ability to alter IAV transmission in our system. These bacterial sialidases preferentially cleave 2,3-, 2,6-, and 2,8- or only 2,3-linked SA, respectively (42). We found that by colonizing mice with the Spn*nanA^-^nanB^-^* mutant, we completely restored the efficiency of IAV transmission from 32% to 100% (*p*<0.0001) (Fig. 5B middle). We then tested the single sialidase mutants (Spn*nanA-* and Spn*nanB-*) and found that the presence of NanA (via Spn*nanB-* colonization) was sufficient to limit IAV acquisition by contacts by 50% (*p*<0.001). Notably, there was no correlation between the colonization density of the bacterial mutants and their effect on shedding or transmission.

To determine the role of sialidases in general in IAV acquisition, infant index and contact mice were treated IN twice daily with *Vibrio cholerae* neuraminidase (VCNA), which cleaves both 2,3- and 2,6-linked sialic acids (43). We found that, like Spn NanA, VCNA treatment was sufficient to decrease IAV acquisition (71.4%) and inhibit shedding by the contacts (Fig. 5B right). Together, these observations suggest that sialidase activity from colonizing bacteria has the capacity to inhibit IAV acquisition in the URT, specifically via its cleavage of 2,3- and 2,6-linked SA.

## DISCUSSION

The inability to study in detail the host’s role in IAV transmission has been a major drawback of the ferret and guinea pig animal models, and has limited our current understanding of IAV contagion (16-19). Herein we established an efficient and tractable infant mouse IAV transmission model with the goal of utilizing the extensive resources of mouse biology, to explore the role that host factors, immune pathways, and the URT flora play in IAV transmission.

Our study corroborated previous findings that infant mice support efficient and consistent IAV transmission (20, 25), and document an age-dependent effect on the efficiency of transmission, highlighting inefficient transmission in adult mice. This suggests an inherent quality of younger mice (i.e. less mobility allowing closer contact, suckling, presence/absence of a host factor, microbiota composition, immunodefficient or developmental status) which facilitates the shedding and transmissibility of IAV. In humans, young age correlates with increased IAV nasopharyngeal shedding (44) and longer duration of shedding (45), increasing the potential for transmission in this age group. Although it has been shown in the study of Edenborough (23) that 56-day old adult mice support the transmission of A/X-31 and A/Udorn/307/72 (H3N2) viruses at 100% efficiency in BALB/c mice, we were unable to observe comparable efficiency in transmission using our A/X-31 virus in C57BL/6J mice older than 28-days of age. The work of Lowen (22) also failed to observe IAV transmission in adult BALB/c mice, further supporting an inconsistent transmission phenotype observed in adult mice which has limited its utilization as an IAV transmission model. Notably, we are the first to demonstrate that infant mice support efficient IBV transmission, which contrasts with the inefficient IBV transmission previously reported in adult mice (32).

Although not evaluated in this study, the difference in mouse strains could also affect the irregular success of IAV transmission in adult mice. One mouse strain, C57BL/6J, has been tested in infant mice and demonstrated 100% transmission efficiency in two independent studies (present study and (25)). In contrast, several mouse strains have been tested for IAV transmission among adult mice, with variable efficiencies among studies (0 to 100%) despite using the similar virus strains. They include BALB/c (22, 23, 26, 28), C57BL/6J (present study), *Mx1*-competent C57BL/6J (24), Swiss Webster (20, 26, 46), New Colony Swiss (47), Manor Farms (MF-1) (32), DBA/2J (26), and Kunming (27). We learned from these studies that, host traits (mouse age, strain, microbiota composition) all contribute to the infectivity and susceptibility of the murine species to IAV, and the host’s contribution to transmission should be explored further using an efficient and tractable model of human disease.

Several studies have demonstrated that virus strain is an important determinant of IAV transmission in mice (20, 23, 32, 46, 48). Like more recent studies (23, 48), we highlighted the increased efficiency of transmission of A/H3N2 over A/H1N1 viruses. In addition, A/H2N2 viruses (not tested here) have also shown to have increased transmission efficiency in mice over A/H1N1 viruses (32, 46). Although we have not evaluated specific viral moieties that confer a transmissible phenotype, the viral HA has been demonstrated to play a role in transmission in mice (23). In addition, we postulate that the activity of some NA in combination with specific HA favors viral release from the nasal epithelium which allows viral shedding and transmission in mice. Thus together, our data highlights that both host and viral-specific features are important to consider to understand the requirements for IAV transmissibility.

We demonstrated that free virus particles present in secretions of mice, and not replicating virus in the URT, correlated with IAV transmission efficiency. A similar observation was also reported by Schulman (32), Edenborough (23), and Carrat (31) demonstrating that transmissibility was associated with greater shedding of virus in index mice, higher viral titers in the saliva of index mice, and shedding of virus from infected humans, respectively. These studies supported our conclusion that viruses which replicate in the URT without having the ability to exit the host (via shedding) cannot be transmitted efficiently. Additionally, studies by Milton (44) suggested that URT symptoms was associated with nasopharyngeal shedding in humans, and that coughing was not necessary for the spread of infectious virus. This helps explains how mice, which lack the cough reflex, can still produce and shed infectious virions. This emphasizes an important future role of the infant mouse IAV transmission model as a tool to study viral shedding as a surrogate marker of IAV contagion.

Two host traits have been identified in this study that influence IAV transmission: IAV-specific immunoglobulin and the URT microbiota. The passive transfer of maternal immunity is transferred via the placenta pre-natally in an IgG-dependent manner (49, 50), or via breastmilk post-natally mediated by several factors: immunoglobulins, leukocytes, and antimicrobial/anti-inflammatory factors (51-53). Our data recapitulates the value of maternal-infant transfer of IAV-specific IgG as a correlate of infant immunity, by demonstrating a significant inhibitory effect on viral shedding and transmission of infant mice after experimental (inoculation) and natural infection (via transmission). Our experiments also demonstrate that IAV-specific serum IgG is predominantly transferred via breastfeeding in mice, as previously reported (53, 54). The concept of maternal serum IgG passage via breastmilk in mice has not often been recognized, even though it has been shown to occur (53, 55-57). IgG can be synthesized locally in the mammary gland, transferred across the mammary gland epithelium, and subsequently transported from the infant gut to the circulation via neonatal Fc receptors (FcRn) expressed in the proximal intestine (58-60). Although this mechanism of maternal IgG acquisition by infants has not yet been correlated in humans, presence of FcRn in the human intestine has been confirmed (61, 62). Our study does not address the contribution of secretory IgA, which is known to be the most abundant immunoglobulin in breastmilk. Yet, our data suggest that adequate amounts of IAV-specific IgG, which is known to wane within 8 weeks of birth in infants (63), maybe necessary to maintain anti-IAV immunity in the URT and limit IAV transmission in infants, given that at this young age, infants don’t have a fully functional adaptive immune response.

In addition to humoral immunity, our study identified an inhibitory role for the common URT colonizer, Spn, at the step of viral acquisition during transmission of IAV. This phenomenon has never been previously observed, although there has been some evidence suggesting that the preceding colonization of Spn reduces IAV infection (25, 64). Notably, the study by McCullers (64) showed that preceding colonization with Spn protected mice from mortality after IAV challenge, whereas the reverse process: prior infection with IAV with subsequent challenge with Spn yielded the opposite effect. This implied that the timing of pathogen encounter mattered, and the composition of the host microbiota may serve a “prophylactic-like” protective effect. Although no studies have evaluated the role of the respiratory microbiota on IAV transmission, we hypothesize that the differences between the transmissibility of different IAV strains and susceptibility of different populations (infants vs adults) to IAV may be due to a combined effect of the virus’s ability to release from SA and exit the host via shedding, and the susceptibility to viral acquisition by contact hosts based on the composition of their URT microbiota. Our work provides proof-of-principle and highlights the amenability of the infant mouse model as a tool to understand the complex dynamics of virus and host, and their combined effect in IAV transmission.

Lastly, we demonstrate the role of Spn sialidases, NanA and NanB, in antagonizing the acquisition and shedding of IAV by contact mice. We hypothesize that bacteria-driven de-sialylation of the host’s URT glycoproteins for use as nutrient (41), may deplete SA residues necessary for IAV adhesion and infection, thus limiting virus susceptibility, and hence acquisition. Notably, the antagonistic effect of bacterial sialidases on IAV shedding of the index group is not statistically different from uncolonized controls. This is analogous to the clinical effects of NI, whereby oseltamivir treatment of index cases alone has not been shown to reduce viral shedding (10, 11). Only when treatment of both index and naïve contacts were partaken (as in post-exposure prophylaxis), has the effects of NI been effective at preventing acquisition of infection among the contact group (12, 13). Because SA is the primary recognition moiety for many viral respiratory pathogens, the concept of utilizing bacterial sialidases as a broad antiviral agent is currently being explored in humans, although its effect on transmission has not yet been evaluated (65-72).

While the advantages of using murine models are evident, these can also be drawbacks. Humans are genetically diverse, live in complex environments, and have been exposed to a myriad of pathogens, all of which can affect transmissibility and susceptibility to IAV, therefore findings generated in animal models of human disease should always be cautiously interpreted. Nevertheless, studying the complexities of IAV transmission biology in a tractable animal model such as infant mice, will allow intricate and sophisticated investigations, which will further our understanding of IAV contagion that may translate into better vaccines and therapeutics.

## MATERIALS AND METHODS

### Mice

C57BL/6J mice (Jackson Laboratories, ME) were maintained and bred in a conventional animal facility. Pups were housed with their mother for the duration of all experiments. Animal studies were conducted in accordance with the *Guide for the Care and Use of Laboratory Animals* (73), and approved by the *Institutional Animal Care and Use Committee* of NYU Langone Health (Assurance Number A3317-01). All procedures were in compliance with *Biosafety in Microbiological and Biomedical Laboratories*.

### Cells and viruses

Madin-Darby canine kidney (MDCK) cells were cultured in Dulbecco’s modified Eagle’s medium with 10% fetal bovine serum and 1% penicillin-streptomycin (Gibco).

Viruses: A/X-31 (H3N2) [HA/NA genes from A/Aichi/2/1968 and internal genes from A/Puerto Rico/8/1934], a gift from Jan Erikson (U. Penn), whose sequence has been deposited in GenBank (XXXXXX). The following reagents were obtained through BEI Resources (NIAID, NIH): A/X-47 (H3N2) [HA/NA genes from A/Victoria/3/1975 and internal genes from A/Puerto Rico/8/1934] [NR-3663], A/Hong Kong/1/1968-2 MA 21-2 (H3N2) [NR-28634], A/Puerto Rico/8/1934 (H1N1) V-301-011-000 [NR-3169], A/WSN/1933 (H1N1) [NR-2555], A/Brisbane/59/2007 (H1N1) [NR-12282], A/California/4/2009 (H1N1) [NR-13659], B/Lee/1940 V-302-001-000 [NR-3178]. IAV and IBV were propagated in 8-10-day old embryonated chicken eggs (Charles River, CT) for 2 days at 37°C and 33°C, respectively. All viruses were tittered by standard plaque assay in MDCK cells in the presence of TPCK (tolylsulfonyl phenylalanyl chloromethyl ketone)-treated trypsin (Thermo Scientific) (74). Purified virus for ELISA was prepared by harvesting allantoic fluid from eggs containing virus followed by centrifugation (3,000×*g*, 30min, 4°C) to remove debris. Viruses were pelleted through a 30% sucrose cushion (30% sucrose in NTE buffer [100mM NaCl+10mM Tris-HCl+1mM EDTA], pH 7.4) by ultracentrifugation (83,000×*g*, 2hrs), resuspended in phosphate-buffered saline (PBS), and stored at -80°C.

### Virus infection, shedding, and transmission

Pups in a litter (4-7 days of age) were infected (index) with a 3μl sterile PBS inoculum without general anesthesia (to avoid direct lung inoculation) by IN instillation of 250 PFU of IAV (unless otherwise specified), and returned to the litter at the time of inoculation for the duration of the experiment. Shedding of virus was collected by dipping the nares of each mouse in viral media (PBS+0.3% BSA) daily, and samples evaluated via plaque assay. Intralitter transmission was assessed in littermates (contact) at 4-5 days p.i. (day 10-14 of life). Pups and mother were euthanized by CO_2_ asphyxiation followed by cardiac puncture, the URT was subjected to a retrograde lavage (flushing 300μl PBS from the trachea and collecting through the nares), and samples were used to quantify virus (via plaque assay or qRT-PCR) or Spn density (described below). Ratio of index to contact pups ranged from 1:3-1:4.

Where indicated, pups were IN treated twice daily with 90 μU (3μl inoculum) of *Vibrio cholerae* neuraminidase (VCNA, Sigma-Aldrich).

The MID_50_ was calculated by the Reed and Müench method (75).

### Induction of maternal IAV immunity

Adult female mice were infected IN with 250 PFU of A/X-31 in a 6μl inoculum without anesthesia. Mice were left to recover from infection for 7 days prior to breeding. Litters of immune mothers were used in experiments.

### Bacteria strain construction and culture

A streptomycin-resistant derivative of capsule type-4 isolate TIGR4 (**P2406**) was used in this study and cultured on tryptic-soy (TS)-agar-streptomycin (200μg/ml) plates (40). The *nanA-* knockout strain (**P2508**) was constructed by transforming P2406 with genomic DNA from strain P2082 (76) (MasterPure DNA purification kit, Illumina), and selection on TS-agar-chloramphenicol (2.5μg/ml) plates. The *nanB-* knockout strain (**P2511**) was constructed by amplifying the Janus cassette (77) from genomic DNA of strain P2408 (78), with flanking upstream and downstream regions to the *nanB* gene added via isothermal assembly. Strain P2406 was transformed with the PCR product, and the transformants selected on TS-agar-kanamycin (125μg/ml) plates. The *nanA-nanB-* double knockout strain (**P2545**) was constructed by transforming P2511 with genomic DNA from strain P2508, and transformants selected on TS-agar-chloramphenicol plates.

Spn strains were grown statically in TS broth (BD, NJ) at 37°C to an optical density (OD) 620nm of 1.0. For quantitation, serial dilutions (1:10) of the inoculum or URT lavages were plated on TS-agar-antibiotic selection plates with 100μl catalase (30,000 U/ml, Worthington Biochemical) and incubated overnight (37°C, 5% CO_2_). Bacteria were stored in 20% glycerol at −80°C. Colonization of pups was carried out by IN instillation of 10^3^ CFU in 3μl of PBS, 1 or 3 days prior to IAV infection.

### qRT-PCR

Following a retrograde URT lavage with 300µl RLT lysis buffer, RNA was isolated (RNeasy Kit, Qiagen), cDNA was generated (High-Capacity RT kit, Applied Biosystems), and used for quantitative PCR (SYBR Green PCR Master Mix, Applied Biosystems). Results were analyzed using the 2^-ΔΔCT^ method (79) by comparison to GAPDH transcription. Values represent the fold change over uninfected.

### ELISA

Immulon 4 HBX plates (Thermo Scientific) were coated with 5μg/ml purified A/X-31 in coating buffer (0.015M Na_2_CO_3_+0.035M NaHCO_3_, 50μl/well), and incubated overnight, 4°C. After three washes with PBS-T (PBS+0.1% Tween 20, 100μl/well), plates were incubated with blocking solution (BS) (PBS-T+0.5% milk powder+3% goat serum [ThermoFisher], 1hr, 20°C). BS was discarded, mice sera were diluted to a starting concentration of 1:100, then serially diluted 1:2 in BS (100μl/well), and incubated (2hr, 20°C). Three washes with PBS-T was done prior to adding secondary antibody (horseradish peroxidase [HRP]-labeled anti-mouse IgG [whole Ab], GE Healthcare or alkaline phosphatase [AP]-labeled anti-mouse IgA [α chain], Sigma) diluted in BS (1:3000, 50μl/well). After incubation (1hr, 20°C) and three washes with PBS-T, plates were developed for 10min using 100μl/well SigmaFast OPD (*o*-phenylenediamine dihydrochloride [Sigma] and stopped with 3M HCl (50μl/well) or developed for 1-18hr using pNPP (p-nitrophenyl phosphate) [KPL];). Plates were read at OD490nm for the OPD substrate or 405nm for the AP substrate. The endpoint titers were determined by calculating the dilution at which the absorbance is equal to 0.1. The geometric mean titers (GMT) were calculated from the reciprocal of the endpoint titers.

### Statistical analysis

GraphPad Prism 7 software was used for all statistical analyses. Unless otherwise noted, data were analyzed using the Mann-Whitney *U* test to compare two groups, and the Kruskal-Wallis test with Dunn’s post-analysis for multiple group comparisons.

### Data Availability

The authors confirm that data will be made publicly available upon publication upon request, without restriction.

## ACKNOWLEDGMENTS

We thank Elodie Guedin (NYU) for the sequencing of A/X-31. MBO was supported by the NYU Division of Infectious Diseases, Department of Medicine Silverman Scholarship, and Physician Scientist Training Program Award; SB was supported by the NYU School of Medicine and National Institute of Diabetes & Digestive & Kidney Diseases fellowship. This project was supported by grants from the U.S. Public Health Service to JNW (AI038446 and AI105168).

## FIGURE LEGENDS

**Figure S1. Window of IAV transmission.**

As Figure 1C, the window of transmission was evaluated by separating index and contact pups for a defined period prior to contact. After infection of index pups, uninfected contact pups were housed apart (in a separate cage) for additional timepoints, including 24, 48, and 72 hours prior to cohousing with infected index for 5 days. Transmission to contact pups was evaluated via plaque assay from retrograde URT lavages. URT titers are represented by a box plot extending from the minimum to maximum values for each data set. Each symbol represent the titer measured from a single pup with median values indicated by a line within the box. Index and contact pups are shown in black and red symbols, respectively. URT=upper respiratory tract.

**Figure S2. Mode of transmission is via direct pup-to-pup contact.**

Upper panels represent schematic for each experimental condition. The mother or cage contents were evaluated as possible sources of transmission (fomites). (A) All infants in one litter (4-7 days of age) were infected IN with 250 PFU of A/X-31 (index) while the second litter in a separate cage were left uninfected (contact). The mothers from the infected and uninfected cage were exchanged daily without disturbing the pups or cage contents. After 5 days, transmission to contact litter was evaluated via plaque assay from retrograde URT lavages. (B) Index pups in a litter were infected and kept separated from an uninfected contact litter as described above. Cages and cage contents (bedding) from infected and uninfected litters were exchanged daily. After 5 days, transmission to contact litter was evaluated via plaque assay from retrograde URT lavages. (C) Index pups are infected IN with 250 PFU of A/X-31, and cohoused with uninfected littermates (contact) for 5 days prior to evaluating for transmission. Transmission to pups and mother were evaluated via plaque assay from retrograde URT lavages. (D) Mother was infected IN with 250 PFU of A/X-31 and placed back with her uninfected litter. After 5 days, transmission to pups was evaluated via plaque assay from retrograde URT lavages.

URT titers are represented by a box plot extending from the minimum to maximum values for each data set. Each symbol represent the titer measured from a single pup with median values indicated by a line within the box. Index and contact pups are shown in black and red symbols, respectively. IAV=Influenza A virus, URT=upper respiratory tract, PFU=plaque forming unit, LOD=Limit of detection.

**Figure S3. Timing of IAV shedding corresponds to timing of transmission.**

Overlay of shedding data (Fig. 2) (black and red symbols) with transmission data (Fig. 1D). Percent transmission was calculated for each day from raw data in Fig. 1D, and is indicated by blue symbols connected by a blue line. Area under the curve corresponds to the proportion of contact pups that have likely acquired IAV infection. IAV=Influenza A virus, PFU=plaque forming unit, LOD=Limit of detection.

**Figure S4. Correlation analyses of influenza virus transmission.**

(A) Mean URT titers (black) and mean shedding titers (green) from index pups were compared to transmission efficiency in contact pups. (B) Index pup shedding titers were compared among various influenza virus strains. Median values are indicated. Threshold of transmission is displayed with dashed line, and was calculated using best-fit linear regression line equation in (A) when transmission X=33.3%. This represents the minimum level of shedding likely to result in transmission of 1 out of 3 contact pups. Each symbol represent the shedding titer measured from a single pup. Shedding titers shown represent at least 2 independent experiments. Transmissible virus and non-transmissible viruses are shown in green and gray symbols, respectively. (C) Age of mice (orange) was compared to transmission efficiency in contact mice. (D) Mean URT titers from index adult mice of 35 days of age (black) and transmission efficiency in contact adult mice of 35 days of age (blue) were compared to index mice inoculum virus titer.

Pearson correlation (*r*) was calculated for each data set. Best fit linear regression curves were fitted on data sets with adequate correlation and goodness of fit (*R^2^*) was calculated. Significance of *r* and *R^2^* were calculated automatically by Graphpad Prism 7 software, * *p*<0.05.

**Figure S5. Serum IgA levels does not correspond to inhibition of shedding.**

As per Fig. 3, serum from mother and pups were obtained at the time of sacrifice. Samples from individual mice were evaluated for IAV-specific IgA by ELISA. Assay controls include serum-deficient PBS samples (Neg) and normal mouse serum (N Serum). IgA geometric mean titers (GMT) are represented by a box plot extending from the 25^th^ to 75^th^ percentile for each data set. Whiskers for each box encompasses the minimum to maximum values. Median values are indicated by a line within the box. All panels represent at least two independent experiments. PFU=plaque forming unit, LOD=Limit of detection, GMT=Geometric mean titer.

